# Beyond Single-Metric Assessments: Uncovering Masked Butterfly Declines via Multi-Scalar Analysis in Central Alberta

**DOI:** 10.64898/2026.08.27.747689

**Authors:** Natalia Lifshitz, Delano Lewis

## Abstract

1. This study analyzed 21 years (2000–2025) of butterfly count data from Central Alberta, integrated with intensive 5-year (2021–2025) high-resolution intra-seasonal sampling.
2. Long-term macro-scale analysis revealed a significant decline in Shannon Diversity, a change that remained obscured when relying solely on traditional metrics of species richness and evenness.
3. This diversity decline was primarily driven by the severe, long-term collapse of the native Common Ringlet (*Coenonympha tullia*).
4. Four other dominant species—Cabbage White (Pieris rapae), Clouded Sulphur (*Colias eriphyle*), European Skipper (*Thymelicus lineola*), and Common Wood Nymph (*Cercyonis pegala*)— maintained long-term population stability, though their abundances were significantly constrained by extreme winter minimum temperatures and rapid spring warming
5. High-resolution intra-seasonal analysis (2021–2025) demonstrated that community indices and species-specific abundances were strongly limited by daily weather, particularly wind velocity and temperature.
6. These findings illustrate that while traditional metrics like richness and evenness are fundamental to community ecology, they provide incomplete insights when applied in isolation; they are most effective when utilized as part of a complementary, multi-scalar framework.
7. This study highlights the necessity of coupling multi-decadal historical datasets with high-frequency, fine-scale sampling to accurately identify the mechanisms of community turnover that simpler metrics may overlook.
8. The results underscore the critical importance of standardized citizen science monitoring in quantifying environmental impacts and establishing conservation priorities for terrestrial insect groups.

---

For many years researchers have noticed declines in insect populations across the globe. This has been called the “insect apocalypse” following a landmark study that showed over 75% decline in flying insect biomass in a 27-year period (Hallmann *et al*., 2017). In Canada and the USA, this decline is an alarming 22% in just 20 years (Edwards *et al*., 2025). These declines have been linked to climate change (Forister *et al*., 2021) with some researchers noting that as ranges become variable the health of ecosystems that have lost butterflies become fragile (Gezon *et al*., 2018). Declines in butterfly populations are driven by habitat change (loss, fragmentation, and/or degradation) with anthropogenic climate change having compounding effects (Kocsis & Hufnagel, 2011; Forister *et al*., 2023).

Climate change acts as a primary driver for shifts in the behavior, reproduction, migration, and foraging of wildlife species (Crossley *et al*., 2021; Jones *et al*., 2013). Climate associated environmental stochasticity forces species to adapt to altered niches, frequently pushing their ranges toward the poles and/or higher elevations (Roland & Matter, 2013; Roland & Matter, 2016). Climate change is rapidly shifting weather patterns and eroding the reliability of climate cues used by species to synchronize their life cycles (Abarca *et al*., 2019; Leuenberger *et al*., 2025; Hamon *et al*., 2024). For butterflies, these shifts impact performance, survival, and community structure (Au & Bonebrake, 2019; Habel *et al*., 2019) with some species succumbing to the additional environmental stress because they cannot adapt fast enough; others survive by altering their phenology, the timing of biological events (Crossley *et al*., 2021; Nowicki *et al*., 2008; Wepprich *et al*., 2019). Because species are interconnected, a shift in the phenology or abundance in one species can have impacts at multiple trophic levels, affecting direct and indirect species interactions (Crossley *et al*., 2021; Roland & Matter, 2016). Environmental changes force species to shift ranges (Roland & Matter, 2016; Sattar *et al*., 2021) which in turn can lead to declining genetic diversity in flowering plants (Jones *et al*., 2013).

In some instances, disturbed areas increase habitat for some butterfly species that prefer long grasses (Hof & Svahlin, 2016) resulting in increased population numbers. Cabbage Whites (*Pieris rapae*) and European Skippers (*Thymelicus lineola*) are in greater abundance in more disturbed areas (Riva *et al*., 2018). This can lead to an overall decrease in butterfly species richness even while total abundance remains unaffected. Anthropogenic habitat change should be limited to preserve not just abundance but also diversity (Kocher & Williams, 2000; Casner *et al*., 2014). Ecosystems that appear stable in richness can simultaneously undergo severe structural degradation; this can be masked by simple presence-absence data. To tease out the masking effects created by using only simple metrics, a multi-scalar approach looking at the long-(macro), medium-(meso), as well as short-term (micro) trends is recommended.

Insect populations are driven by stressors operating on three different time scales; macro-scale (decades), meso-scale (month to years), and micro-scale (hours to weeks). Studies using aggregated butterfly count data (macro-scale) and in some areas counting butterflies in the summer months are increasing in popularity (micro-scale), but early-winter weather (meso-scale) is a pivotal driver of butterfly annual population growth, and this is often not considered. For temperate species especially, temperatures in November provide a more accurate prediction of population change than broad climate indices. Species that overwinter as eggs or as pharate larvae face higher risks from extreme cold and unseasonable warm spells in late fall and early winter before they are physiologically prepared (Roland & Matter, 2013; Roland & Matter, 2016). Unseasonably warm winter days can have devastating effects causing 75% to 100% mortality (Abarca *et al*., 2019). The timing of snowmelt and spring warming, which determines the beginning of the thaw cycle, is a critical cue for larvae emerging from hibernation (Roland & Matter, 2016). If larvae emerge before warm weather is consistent or before host plants “leaf out,” they face high mortality (Roland & Matter, 2016; Abarca *et al*., 2019; Leuenberger *et al*., 2025). Late emerging multivoltine species tend to be the most sensitive to inter-annual spring temperature changes (Hamon *et al*., 2024). Additionally, heavy rainfall creates cooler microclimates leading to slower larval development (Abarca *et al*., 2019; Hof & Svahlin, 2016; Pilliod & Rohde, 2016; Chen *et al*., 2019) and limited rainfall during the growing season restricts host plant growth, leading to resource competition and population crashes (Pollard, 1977).

Long-term data is essential for distinguishing between annual population fluctuations and actual temporal trends as we try to understand the drivers of insect declines. A comprehensive study of butterflies across the continental U.S. found that total butterfly abundance plunged 22% between 2000 and 2020 (Wepprich *et al*., 2019). Systematic monitoring over 21 years documented a 33% reduction in total butterfly abundance (Halsch *et al*., 2025), and a 50-year dataset revealed that peak richness days now see 10 fewer species than they did in previous decades (Ries & Oberhauser, 2015). These long-term monitoring studies reveal complex interactions in butterfly communities with winter colonies showing multi-decadal declines while summer counts reveal that more nuanced dynamics may be at play (Larsen *et al*., 2024). Long-term data helps to quantify environmental impacts in the context of the stochasticity of butterfly populations (Halsch *et al*., 2025). But while long-term data is required to detect chronic collapses and the influence of global climatic patterns, it often lacks the resolution to explain them. Conversely, fine scale phenological data can explain immediate weather responses but cannot establish a historical baseline. Identifying the true mechanisms of community turnover and population stochasticity requires coupling multi-decadal historical data with intensive, high-frequency intra-seasonal sampling to capture both the “slow” crashes and the “fast” weather filters. Studying butterfly diversity becomes important given that their diversity correlates with the diversity of other terrestrial insect groups that make up the bulk of global biodiversity (Nowicki *et al*., 2008; Larsen *et al*., 2024). By tracking population shifts in butterflies over decades, monitoring programs can provide the long-term data necessary to quantify environmental impacts and establish conservation priorities (Wepprich *et al*., 2019; Larsen *et al*., 2024).

Simple count data are foundational for calculating species richness (S), a measure of species occupancy, and species evenness (J), which quantifies the distribution of abundance among those species. These traditional metrics are widely utilized as standard descriptors of ecosystem health (MacDonald et al., 2017; Bollarapu et al., 2024). However, when interpreted in isolation, richness and evenness may lack the sensitivity required to detect nuanced temporal shifts, particularly in ecosystems where a small number of dominant taxa dictate the overarching community structure. Recent comparative assessments suggest that while Simpson’s (D) and Shannon (H) indices offer different sensitivities with D better detecting dominance patterns and H responding more dynamically to rare species, no single index captures the full complexity of ecological change (Dubey et al., 2025). Rather than viewing these as competing metrics, we propose a complementary approach by integrating traditional diversity indices with high-resolution, multi-scalar analyses. In this way we can move beyond single metric limitations to reveal structural degradation that would otherwise be masked by stable presence-absence data.

Butterfly monitoring through counts is a global model for citizen science, transforming public engagement into a powerful engine for ecological research and helping to generate a huge volume of reliable data (Prudic *et al*., 2017). While some researchers have questioned the reliability of non-professional data, current models show that data from citizen scientists are comparable in quality, accuracy, and reliability to professional efforts (Wepprich *et al*., 2019; Ries & Oberhauser, 2015; Prudic *et al*., 2017). Comparing mass-participation projects like the Big Butterfly Count (BBC) with professional efforts such as done by the United Kingdom Butterfly Monitoring Scheme (UKBMS) show that simple sampling protocols can produce comparable and reliable estimates of species abundance (Prudic *et al*., 2017; Van Strien *et al*., 2017; Sevilleja *et al*., 2020). Organizations such as the North American Butterfly Association (NABA) and digital platforms like eButterfly and iNaturalist aggregate millions of records across entire continents, providing insights into distribution and population sizes that professional scientists could not achieve on their own (Crossley *et al*., 2021; Ries & Oberhauser, 2015). Standardization in counts is necessary; variations of the Pollard walk (Pollard, 1977) are usually used to track species richness and diversity over time. Using such methods, in North America alone NABA facilitates counts at over 400 sites, providing critical data on species distribution, population sizes, and the impacts of climate and habitat change (Halsch *et al*., 2025).

Historically, five butterfly taxa dominate the butterfly community in central Alberta: the Cabbage White (*Pieris rapae*); the Clouded Sulphur (*Colias eriphyle*); the Canadian Tiger Swallowtail (*Papilio canadensis*); the Mourning Cloak (*Nymphalis antiopa*); and the Common Ringlet (*Coenonympha tullia*) (Ryan *et al*., 2019; Flockhart, 2002; Konno, 2023; Teichman *et al*., 2013). The list of dominant species will vary depending on the location, but you often find both widespread species (such as the Cabbage White, the Clouded Sulphur, and the Canadian Tiger Swallowtail), and more localized residents (such as the Ringlet and the Mourning Cloak). The European Skipper (*Thymelicus lineola*), an introduced grass feeder, is historically not in the top 5 species across central Alberta but can be extremely abundant in specific areas. Other butterflies that can be locally abundant include the White Admiral (*Limenitis arthemis rubrofasciata*), the Silvery Blue (*Glaucopsyche lygdamus couperi*), and the Common Wood Nymph (*Glaucopsyche lygdamus couperi*) (Flockhart, 2002).

The non-for-profit organization where this study was conducted has been conducting butterfly counts during an annual event since 2000. In 2021 a more intensive count was initiated where several surveys were done over the summer months of June, July, and August at that location. These counts have revealed that the five most dominant taxa there are the Cabbage White (*Pieris rapae*), the Clouded Sulphur (*Colias eriphyle*), the European Skipper (*Thymelicus lineola*), the Common Ringlet (*Coenonympha tullia*), and the Common Wood Nymph (*Cercyonis pegala*).

The aims of this study are to: (i) assess multi-decadal shifts in community-level diversity (H, S, J); (ii) isolate the long-term temporal trends from macro-climate noise over 21 years; and (iii) use the intensive 5-year dataset to map the fine-scale, intra-seasonal ecological fingerprints of these dominant species.

## Results

### Long-Term Drivers of Community Structure and Population Dynamics

Over the 21-year macro-temporal scale, Species Evenness (J) exhibited no significant directional trend, remaining relatively stable despite the collapse of the Ringlet (Table 1, Fig. 1). However, our fine-scale analysis of the 5-year dataset reveals that overall evenness undergoes a significant seasonal decline, as the community becomes increasingly dominated by a few abundant taxa like the Cabbage White toward the end of the summer. While evenness does not decline with time, there is a predictable intra-seasonal “structural shift” where dominance patterns intensify as the flight season progresses. Shannon Diversity (H) experienced a significant long-term decline (Standardized β = −0.019, 95% CI: −0.035 to −0.003; Relative Variable Importance [RVI] = 0.85; Fig. 2). Furthermore, H was positively impacted by the temperature on the day of the survey (standardized β = 0.11, RVI = 0.68. 95% CI [0.016, 0.204]) and Snow Exposure Index (Standardized β = 0.113, RVI = 0.32, 95% CI [0.005, 0.221]) and negatively correlated to rapid spring warming (Spring GDD: β = −0.112, RVI = 1.00). Goodness-of-fit evaluations indicated that the top candidate models for H captured a substantial portion of the historical variance (R^2^m = 0.38–0.56). In contrast, the top model subsets for S (R^2^m = 0.00–0.18) and J (R^2^m = 0.00–0.16) explained considerably less variance, reflecting the retention of the intercept-only null models within their respective equally parsimonious subsets (ΔAICc < 2).

**Figure 1.**
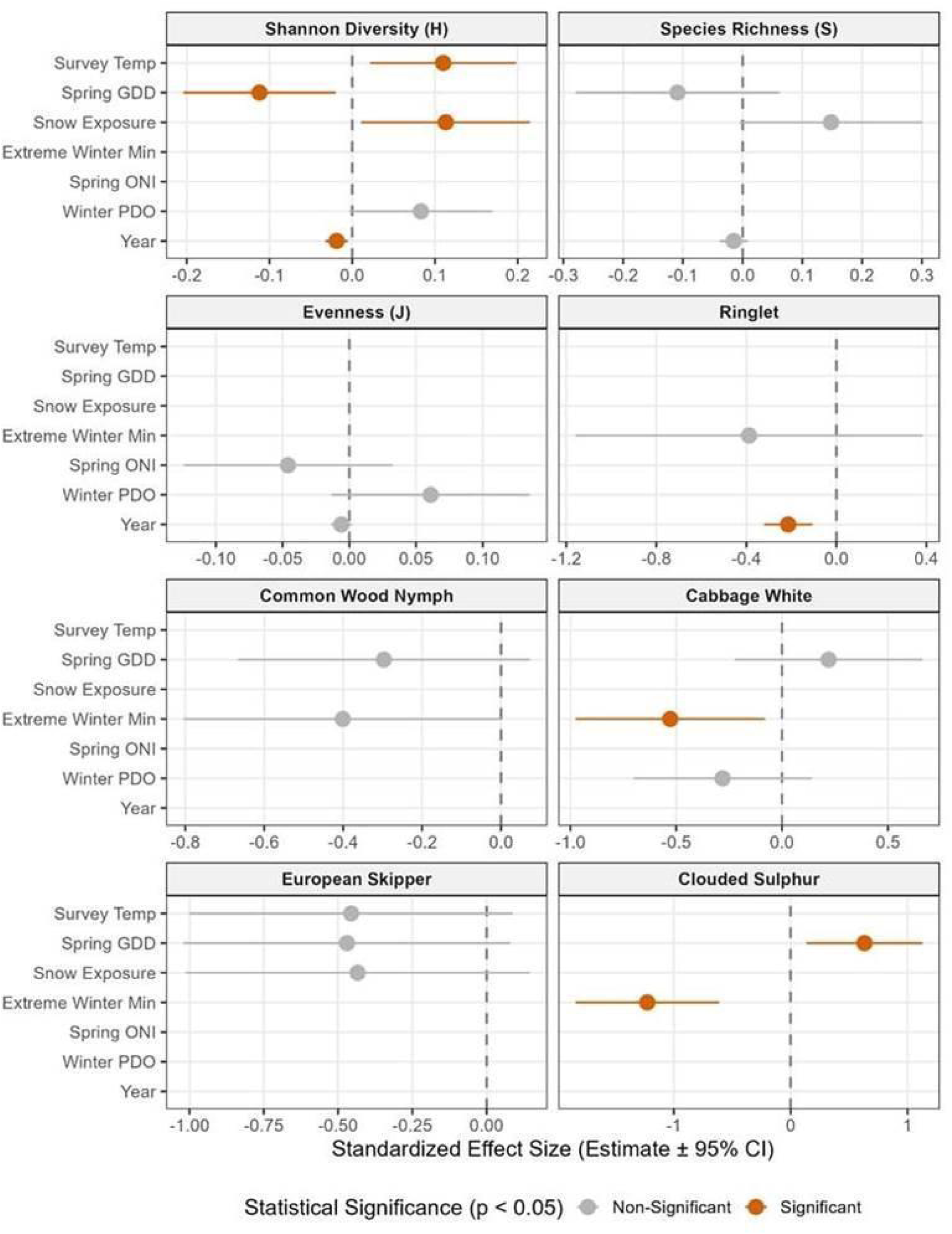
Standardized effect sizes of temporal and climatic drivers on butterfly community structure and dominant species abundances (21-year data). Forest plot displaying conditional model-averaged estimates (points) and 95% confidence intervals (horizontal bars) derived from the most parsimonious generalized linear models (ΔAICc < 2). Predictor variables were centered and standardized prior to analysis to allow for direct comparison of effect sizes across models. The vertical dashed line indicates an effect size of zero (no effect). Highlighted orange points indicate statistically significant predictors (p < 0.05), while gray points represent non-significant variables retained in the top model subset. The plot illustrates the severe, asynchronous long-term decline (Year) in Shannon Diversity (*H*) and the native Ringlet population, contrasting with the temporal stability of the remaining dominant taxa. (Note: Clouded Sulphur estimates represent the single best model, as no other models competed within ΔAICc < 2).

**Figure 2.**
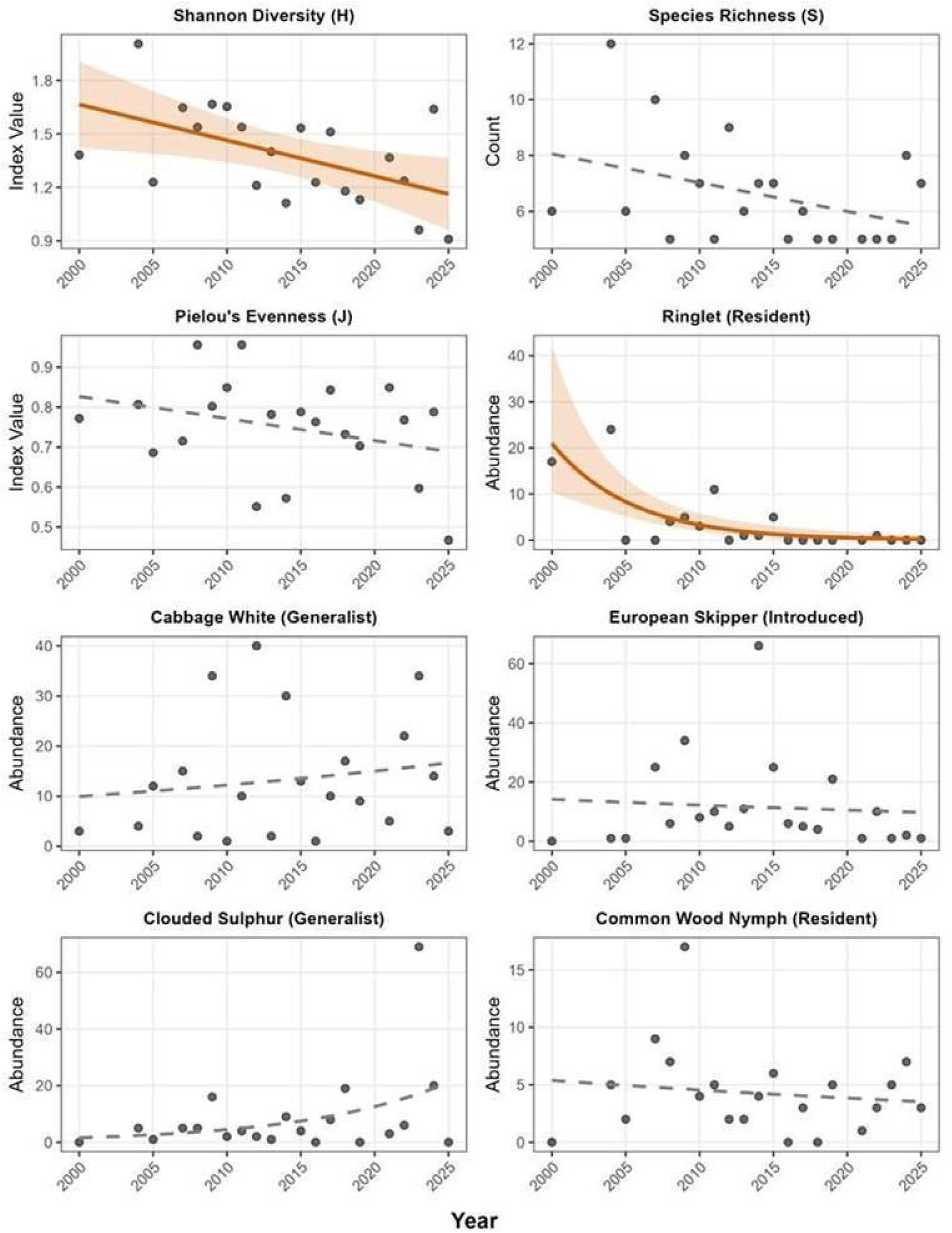
Long-term temporal trends in butterfly community structure and the abundance of dominant taxa (21-year data). Scatter plots display raw annual values for community indices (Shannon Diversity *H*, Species Richness *S*, Pielou’s Evenness *J*) and raw observation counts for the five most historically abundant butterfly species. Trendlines represent fitted model predictions: linear models were applied to community indices, and Negative Binomial generalized linear models were applied to abundance counts to account for overdispersed variance structures. Shaded regions represent 95% confidence intervals. Bold, colored solid lines highlight statistically significant temporal declines (p < 0.05) identified via multi-model inference. Dashed gray lines indicate non-significant temporal trends, representing long-term population stability. The pronounced collapse of the native Ringlet strongly contrasts with the stability of other dominant taxa, illustrating the demographic mechanism driving the concurrent decline in Shannon Diversity.

**Table 1.** Long-Term Drivers of Butterfly Community Structure and Dominant Species Abundance (21-year data) Summary of model-averaged coefficients evaluating the effects of temporal trends and macro-climatic drivers over a 21-year period. Values represent the conditional average across the top model subset (ΔAICc < 2). Predictors are standardized (Z-scored) to allow direct comparison of effect sizes (β). Relative Variable Importance (RVI) represents the sum of Akaike weights (w_i_) for all models containing the predictor. Bold p-values indicate statistical significance (p < 0.05). Clouded Sulphur values represent the single most parsimonious model (ΔAICc = 0.00), hence RVI = 1.00.

| Response variable | Predictor Retained | Standardized $\beta$ | Adj. S.E. | 95% CI | RVI | p-value |
| --- | --- | --- | --- | --- | --- | --- |
| **Community Indices** |  |  |  |  |  |  |
| Shannon Diversity (H) | Spring GDD | -0.112 | 0.050 | [-0.210, -0.014] | 1.00 | **0.026** |
|  | Year | -0.019 | 0.008 | [-0.035, -0.003] | 0.85 | **0.014** |
|  | Survey Temp | 0.110 | 0.048 | [0.016, 0.204] | 0.68 | **0.023** |
|  | Winter PDO | 0.083 | 0.047 | [-0.009, 0.175] | 0.36 | **0.081** |
|  | Snow Exposure Index | 0.113 | 0.055 | [0.005, 0.221] | 0.32 | **0.041** |
| Species Richness (S) | Snow Exposure Index | 0.148 | 0.083 | [-0.015, 0.311] | 0.52 | 0.075 |
|  | Spring GDD | -0.109 | 0.093 | [-0.291, 0.073] | 0.32 | 0.241 |
|  | Year | -0.015 | 0.013 | [-0.040, 0.010] | 0.14 | 0.241 |
| Pielou's Evenness (J) | Spring ONI | -0.046 | 0.040 | [-0.124, 0.032] | 0.32 | 0.247 |
|  | Year | -0.006 | 0.004 | [-0.014, 0.002] | 0.29 | 0.179 |
|  | Winter PDO | 0.061 | 0.038 | [-0.013, 0.135] | 0.16 | 0.110 |
| **Species Abundance** |  |  |  |  |  |  |
| Ringlet | Year | -0.214 | 0.055 | [-0.322, -0.106] | 1.00 | **<0.001** |
|  | Extreme Winter Min | -0.388 | 0.393 | [-1.158, 0.382] | 0.29 | 0.324 |
| Cabbage White | Extreme Winter Min | -0.528 | 0.228 | [-0.975, -0.081] | 0.51 | **0.021** |
|  | Winter PDO | -0.281 | 0.215 | [-0.702, 0.140] | 0.21 | 0.191 |
|  | Spring GDD | 0.220 | 0.226 | [-0.223, 0.663] | 0.14 | 0.330 |
| European Skipper | Survey Temp | -0.456 | 0.278 | [-1.001, 0.089] | 0.37 | 0.101 |
|  | Snow Exposure Index | -0.434 | 0.296 | [-1.014, 0.146] | 0.31 | 0.143 |
|  | Spring GDD | -0.470 | 0.281 | [-1.021, 0.081] | 0.15 | 0.094 |
| Common Wood Nymph | Extreme Winter Min | -0.401 | 0.206 | [-0.805, 0.003] | 0.43 | 0.051 |
|  | Spring GDD | -0.297 | 0.189 | [-0.667, 0.073] | 0.28 | 0.117 |
| Clouded Sulphur | Extreme Winter Min | -1.227 | 0.313 | [-1.840, -0.614] | 1.00 | **0.001** |
|  | Spring GDD | 0.632 | 0.253 | [0.136, 1.128] | 1.00 | **0.022** |

The five dominant species showed species-specific sensitivity to large-scale climate and local weather. The Ringlet exhibited a strong, highly significant temporal decline (β = −0.214, RVI = 1.00; Table 1, Fig 2), with its top candidate models explaining nearly 60% of the population variance (R^2^m = 0.59). In contrast, the remaining dominant taxa maintained long-term population stability (Fig. 1, Fig. 2). Instead, their abundance was dictated primarily by macro-climatic variables. The single most parsimonious model for the Clouded Sulphur (ΔAICc = 0.00) revealed it was highly sensitive to extreme winter minimums (β = −1.227) and rapid spring warming (spring GDD; β = 0.632), similarly capturing a high degree of historical variance (R^2^m = 0.58). The Cabbage White was significantly affected by the extreme winter minimum (β = −0.528, p = 0.021) and for the remaining stable species (European Skipper and Common Wood Nymph), the variance explained by the top macro-climatic models was more moderate, with R^2^m values ranging from 0.00 to 0.29.

### Intra-Seasonal High-Resolution Dynamics (2021–2025)

High-resolution modeling of the 5-year dataset (Table 2, Fig. 3) revealed that immediate day-of weather conditions and phenology tightly govern community detectability and intra-seasonal flight activity. Both Shannon Diversity (H) and Species Richness (S) were significantly enhanced by optimal thermal windows (Day-of Temp RVI = 0.74 and 1.00, respectively) and strongly suppressed by wind velocity (Day-of Wind RVI = 1.00 for both indices). Additionally, a strong negative effect of Julian Day across H, S, and J (all RVI = 1.00) indicates a general concentration of overarching community richness and evenness toward the earlier half of the mid-summer survey window. Goodness-of-fit for top candidate models indicated that fixed and random effects explained approximately 20–22% of the variance for H (R^2^m = 0.20–0.22; R^2^c = 0.20–0.22) and S (R^2^m = 0.21–0.22; R^2^c = 0.21–0.22), and between 75–81% of the variance for J (R^2^m = 0.75–0.81; R^2^c = 0.75–0.81).

**Figure 3.**
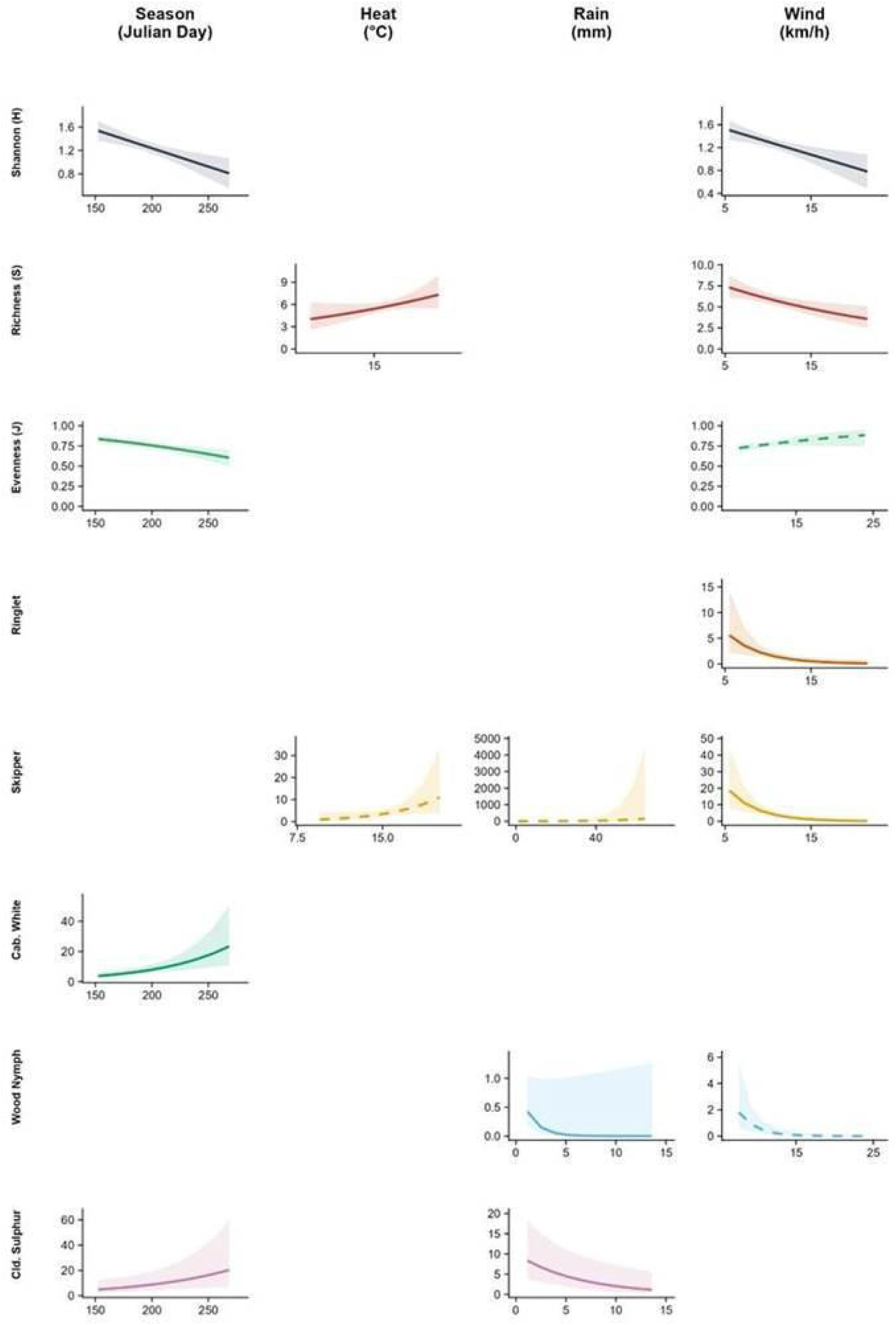
Ecological Fingerprint Matrix of Community Indices and Dominant Butterfly Species (2021– 2025). Each row represents a response variable, including community diversity (Shannon *H*), richness (*S*), evenness (Pielou’s *J*), and the abundance of the five most common species. Columns correspond to primary weather categories: Seasonality (Julian Day), Heat (°C), Precipitation (mm), and Wind Speed (km/h). Individual panels display predicted values (solid or dashed lines) with 95% confidence intervals (shaded areas) derived from Generalized Linear Mixed Models (GLMMs). Solid lines represent responses to instantaneous weather at the time of sampling, while dashed lines indicate responses to 3-day lagged weather variables (e.g., cumulative rain or average temperature prior to the survey). Empty panels indicate variables not retained in the top-performing model subset (ΔAICc < 2). X-axes are scaled to the observed range of weather conditions in the field to prevent over-extrapolation.

**Table 2.** Intra-Seasonal Weather and Phenological Drivers of Butterfly Dynamics (5-Year High-Resolution Dataset) Summary of model-averaged coefficients evaluating the effects of day-of and lagged (3-day) weather conditions, alongside survey phenology (Julian Day), on community indices and species abundance. Values represent the conditional average across the top model subset (ΔAICc < 2). Predictors are standardized (Z-scored) to allow direct comparison of effect sizes (β). Relative Variable Importance (RVI) represents the sum of Akaike weights (w_i_) for all models containing the predictor. Bold p-values indicate statistical significance (p < 0.05).

| Response Variable | Predictor Retained | Standardized $\beta$ | Adj. SE | 95% CI | RVI | p-value |
| --- | --- | --- | --- | --- | --- | --- |
| **Community Indices** |  |  |  |  |  |  |
| Shannon Diversity (H) | Julian Day | -0.177 | 0.050 | [-0.275, -0.079] | 1.00 | **<0.001** |
|  | Wind (Day of) | -0.156 | 0.049 | [-0.252, -0.060] | 1.00 | **0.001** |
|  | Temp (Day of) | 0.113 | 0.049 | [0.017, 0.209] | 0.74 | **0.021** |
|  | Temp (3-Day Avg) | 0.095 | 0.049 | [-0.001, 0.191] | 0.26 | 0.052 |
|  | Rain (3-Day Sum) | 0.040 | 0.048 | [-0.054, 0.134] | 0.22 | 0.404 |
| Species Richness (S) | Julian Day | -0.115 | 0.054 | [-0.221, -0.009] | 1.00 | **0.034** |
|  | Temp (Day of) | 0.163 | 0.052 | [0.061, 0.265] | 1.00 | **0.002** |
|  | Wind (Day of) | -0.163 | 0.054 | [-0.269, -0.057] | 1.00 | **0.003** |
|  | Rain (3-Day Sum) | 0.039 | 0.050 | [-0.059, 0.137] | 0.29 | 0.439 |
| Pielou's Evenness (J) | Julian Day | -0.308 | 0.087 | [-0.479, -0.137] | 1.00 | **<0.001** |
|  | Wind (3-Day Avg) | 0.141 | 0.099 | [-0.053, 0.335] | 0.45 | 0.154 |
|  | Temp (3-Day Avg) | -0.134 | 0.092 | [-0.314, 0.046] | 0.36 | 0.145 |
|  | Rain (Day of) | 0.101 | 0.094 | [-0.083, 0.285] | 0.18 | 0.280 |
|  | Temp (Day of) | -0.112 | 0.096 | [-0.300, 0.076] | 0.16 | 0.244 |
|  | Wind (Day of) | -0.082 | 0.094 | [-0.266, 0.102] | 0.13 | 0.384 |
|  | Rain (3-Day Sum) | -0.087 | 0.106 | [-0.295, 0.121] | 0.06 | 0.411 |
| **Species Abundance** |  |  |  |  |  |  |
| Ringlet | Wind (Day of) | -0.875 | 0.315 | [-1.492, -0.258] | 1.00 | **0.005** |
|  | Julian Day | -0.317 | 0.287 | [-0.880, 0.246] | 0.34 | 0.269 |
|  | Temp (Day of) | 0.332 | 0.219 | [-0.097, 0.761] | 0.32 | 0.130 |
|  | Temp (3-Day Avg) | 0.332 | 0.230 | [-0.119, 0.783] | 0.28 | 0.149 |
|  | Rain (3-Day Sum) | -0.172 | 0.289 | [-0.738, 0.394] | 0.08 | 0.552 |
| Cabbage White | Julian Day | 0.518 | 0.147 | [0.230, 0.806] | 1.00 | **<0.001** |
|  | Temp (Day of) | 0.348 | 0.130 | [0.093, 0.603] | 0.84 | **0.007** |
|  | Wind (Day of) | -0.239 | 0.135 | [-0.504, 0.026] | 0.58 | 0.077 |
|  | Rain (Day of) | -0.171 | 0.142 | [-0.449, 0.107] | 0.34 | 0.228 |
|  | Wind (3-Day Avg) | -0.310 | 0.123 | [-0.551, -0.069] | 0.17 | **0.012** |
| European Skipper | Temp (3-Day Avg) | 1.113 | 0.290 | [0.545, 1.681] | 1.00 | **<0.001** |
|  | Wind (Day of) | -0.979 | 0.272 | [-1.512, -0.446] | 1.00 | **<0.001** |
|  | Rain (3-Day Sum) | 0.356 | 0.367 | [-0.363, 1.075] | 0.34 | 0.333 |
| Common Wood Nymph | Rain (Day of) | -2.804 | 1.296 | [-5.344, -0.264] | 1.00 | **0.030** |
|  | Wind (3-Day Avg) | -1.121 | 0.362 | [-1.831, -0.411] | 1.00 | **0.002** |
|  | Julian Day | 0.476 | 0.403 | [-0.314, 1.266] | 0.30 | 0.238 |
|  | Temp (3-Day Avg) | 0.326 | 0.374 | [-0.407, 1.059] | 0.22 | 0.384 |
| Clouded Sulphur | Julian Day | 0.384 | 0.149 | [0.092, 0.676] | 1.00 | **0.010** |
|  | Rain (Day of) | -0.455 | 0.146 | [-0.741, -0.169] | 1.00 | **0.002** |
|  | Wind (3-Day Avg) | -0.351 | 0.165 | [-0.674, -0.028] | 1.00 | **0.034** |
|  | Julian Day<sup>2</sup> | -0.099 | 0.118 | [-0.330, 0.132] | 0.23 | 0.402 |
|  | Temp (3-Day Avg) | 0.103 | 0.153 | [-0.197, 0.403] | 0.21 | 0.499 |

At the species level, the conditional model averages highlighted highly varied phenological strategies and meteorological thresholds (Fig. 3). The Cabbage White and Clouded Sulphur exhibited strong positive associations with Julian Day (β = 0.518 and 0.384, respectively; Table 2). Conversely, flight activity for the introduced European Skipper was influenced by localized warming trends over the preceding days (3-Day Average Temp: β = 1.113, RVI = 1.00) as well as wind (Day of: β =−0.979, RVI = 1.00). Notably, the residents demonstrated acute sensitivity to immediate atmospheric stressors: the Ringlet was severely suppressed by day-of wind (β = −0.875, RVI = 1.00), while the Common Wood Nymph was negatively impacted by both immediate precipitation (β = −2.804, RVI = 1.00) and lagged 3-day wind conditions (β = −1.121, RVI = 1.00). Across the species-specific top models, the variance explained exclusively by fixed effects (R^2^m) ranged from 0.18 for the Ringlet to 0.70 for the Common Wood Nymph. The total variance explained when incorporating the survey year random effect (R^2^c) remained nearly identical to the marginal variance for the Ringlet (R^2^c = 0.18–0.20) and Common Wood Nymph (R^2^c = 0.68–0.70), but increased notably for the Clouded Sulphur (R^2^m = 0.22; R^2^c = 0.51–0.53) and Cabbage White (R^2^m = 0.34–0.39; R^2^c = 0.43–0.46)].

## Discussion

### Long-term Diversity Decline and Dominant Species (Macro-scale)

Over the 21-year study period, the butterfly community exhibited a significant structural shift; diversity declined, but this was not driven by localized extirpations, since species richness remained stable across the same period and evenness showed no significant directional trend. Species-specific abundance modeling of the five dominant taxa revealed a clear mechanistic driver for the overarching shift in community evenness. Species richness and evenness showed low historical variance (between 0.00 and 0.18 and 0.00 and 0.16 respectively) which suggests that localized, stochastic events are better predictors for these metrics than multi-decadal climatic factors. The Ringlet, a historically dominant native resident, underwent a severe and highly significant population collapse over the 21-year timeline. The goodness-of-fit analysis revealed that more than half of the historical variance could be explained using a few climatic and temporal predictors - this is rare in datasets. Our study found that overall temporal shifts were predictive of the diversity declines in the Ringlet which reflects a study on UK butterflies which found that extreme winter minimums was a key predictor for *Aphantopus hyperantus* populations (McDermott Long *et al*., 2017). Other researchers have found that macro-climatic variables are “vital predictors” of butterfly population dynamics and extinction risks (Li *et al*., 2023). Pinkert *et al*. (2025) found that diversity in butterfly populations, specifically those containing mountain specialists, is exceptionally threatened by the erosion of extreme winter minimums which they identify as the primary driver of diversity declines.

During model validation several high-leverage data points were examined to see if they were influencing (or skewing) the long-term data trends. We found that these points consistently corresponded with years that had extreme climatic events and so we retained them rather than excluded them since these climatic events, such as severe winter minimums and rapid spring warming were already a part of our model. This can be seen in years that have disproportionate population crashes corresponding to periods of extreme thermal instability or record-breaking cold spells. This demonstrates that there is sensitivity of these populations to environmental stochasticity and not mere sampling biases or data artifacts.

### Effects of Winter Temperatures and Seasonal Changes in Evenness (Meso-scale)

The asynchronous population dynamics observed where the resident Ringlet experienced acute collapse while another resident, the Common Wood-nymph and more widespread species such as the Cabbage White, the Clouded Sulphur, the European Skipper, maintain stable baselines agree with previous research that shows that decreases in diversity are frequently driven by the disappearance of habitat specialists, likely to be residents, rather than the dominant generalists which are often more widespread. Species with high habitat specificity show higher sensitivity to urbanization and human disturbance than mobile generalists (Kuussaari *et al*., 2021). Research also shows that the more common species are less likely to drive declines in species richness because they are still being encountered even after moderate population declines, while less common species are more likely to disappear completely from survey records (Edwards *et al*., 2025).

In contrast to the Ringlet, the remaining four most abundant species, comprising two highly mobile and more widespread species, an introduced grass-feeder, and another native resident, demonstrated long-term temporal stability. Rather than time-driven declines, the abundances of these stable survivors were constrained primarily by localized weather filters. Extreme winter minimum temperatures had a strong influence in both the Cabbage White and the Clouded Sulphur and a marginal influence in the Common Wood Nymph. This indicates that colder, harsher winter extremes cause higher summer abundances for these taxa. Additionally, rapid spring warming significantly bolstered Clouded Sulphur, possibly through accelerated development, earlier spring activity, or increased voltinism, which has been observed in other butterflies (Ragonese *et al*., 2024; Vitasse *et al*., 2021). The goodness-of-fit models reveal that highly mobile, and more widespread multivoltine species, such as the Cabbage White and Clouded Sulphur, showed a boom-and-bust pattern with massive year-to-year population fluctuations layered on top of immediate daily responses to weather; this leads to long term population stability in these butterflies (de Palma *et al*., 2016; Hussain *et al*., 2018). By adding meso-scale metrics to the larger macro-scale trends we finally start to see better resolution and uncover mechanisms of community level changes that are occurring.

### Short-term Seasonal Changes and the Responses of Dominant Species (Micro-scale)

Looking closer into detailed, intra-seasonal patterns, we found that diversity shows a strong seasonal decline, driven by Julian Day. Evenness also declines through the season, indicating that the butterfly community in our study area becomes increasingly dominated by a few abundant taxa as the season progresses. This is supported by research that shows the Cabbage White starting the season with a relatively small spring generation but shows drastic population booms in consequent generations (von Schmalensee *et al*., 2023). Fine-scale community dynamics were strongly limited by count-day wind speed, which acted as a primary filter for Species Richness (S) and Shannon Diversity (H) by reducing the number of observed species. In contrast, Species Evenness (J) was not significantly impacted by either count-day wind or the 3-day wind average. This suggests that while wind conditions reduce the overall detectability of the community, they do not significantly alter the proportional dominance of the species that remain active and visible. Previous research found that environmental variability (including wind) can reduce the dominance of the most competitive species and promote a more diverse community structure (Ulrich *et al*., 2016). Species richness was primarily limited by count-day wind conditions, with higher wind speeds causing a significant reduction in the number of observed species; this reflects studies on birds and flying foxes that show similar results (Robbins, 1981; Dorrestein *et al*., 2025). Count-day temperature was also a strong positive driver, significantly outperforming the 3-day temperature lag, indicating that richness is driven by current thermal opportunity rather than physiological history. Previous studies suggest that species richness is strongly influenced by current temperatures, often being a dominant driver of observed richness (Mena *et al*., 2025). A seasonal decline in richness was also observed, though this phenological signal was slightly weaker than that observed for Shannon diversity or Evenness. This was likely the result of climate, resource availability, and or physiological constraints. While community-level indices (H, S, and J) showed no significant response to precipitation variables in the high-resolution dataset, immediate weather filters remained critical at the species level. Specifically, day-of rain significantly suppressed the flight activity and detectability of the Clouded Sulphur and the Common Wood Nymph. This suggests that while the abundance of both residents and more widespread species is sensitive to moisture, these effects do not aggregate into a detectable shift in total community diversity or richness during the survey window.

Wind on the day of the count was another predictor of Ringlet number, decreasing counts with increasing wind speeds. Interestingly, the Ringlet showed little response to thermal history or precipitation, suggesting a strategy of opportunistic flight whenever wind conditions permit. Other studies have found that butterfly assemblages were affected by wind speed and that some species respond differently to wind (Wikström *et al*., 2009). The effect of wind on the Ringlet may be explained by behavioral observations of the Ringlet where it often dives and hides at the base of grasses and plants when disturbed; they may be displaying a similar behavior in strong wind. The European Skipper showed a similar, negative relationship with count-day wind. However, unlike the Ringlet, the Skipper’s abundance was strongly positively associated with 3-day lagged temperature but 3-day lagged precipitation was not significantly correlated; this pattern may suggest that current wind limits detection, warm and moist conditions in the preceding days could likely be a trigger for eclosion events thereby increasing the local pool of available adults. This finding agrees with Leston & Koper (2016) and other research who have found similar ecological patterns (Larsen *et al*., 2024; Leston & Koper, 2019; Kuussaari *et al*., 2016). Ringlets, which overwinter as larvae, tend to thrive in damp summers but they do not have the same positive sensitivity to early season moisture as the European Skipper that overwinters as eggs (Larsen *et al*., 2024). The Common Wood Nymph displayed a unique sensitivity to wet and turbulent weather history; it showed a strong negative response to the 3-day wind average, indicating that sustained periods of high wind significantly reduce observable populations, likely by suppressing activity or increasing mortality rates. These findings are reflective of other work on the Common Wood Nymph in Winnipeg, Canada and related species in Finland (Leston & Koper, 2016; Leston & Koper, 2019; Kuussaari *et al*., 2016). The Cabbage White and the Clouded Sulphur both exhibited significant positive seasonal trends, identifying them as late-season species that increase in abundance while the rest of the community declines. Cabbage whites have been identified as having high voltinism making them more sensitive to shifts in temperatures; this gives them a greater ability to vary the numbers of generations they have each year (Hamon *et al*., 2024) and also cumulatively build up population numbers by fall. Konno (2023) has identified the Cabbage White as having an extremely high growth rate which contributes to its ability to rapidly exploit favorable conditions quickly and complete more life cycles than other species; this also contributes to their high numbers as the season progresses. The Clouded Sulphur showed a significant negative response to count-day rain. However, while the linear model captured this precipitation constraint, residual diagnostics indicated complex quantile deviations, likely attributable to the species’ bi-voltine (two-generation) phenology, which could not be fully resolved by a simple linear or quadratic seasonal term. Count-day rain has been shown to reduce adult activity, lower the rate of detection, and contribute to long-term population declines in several species of butterflies with some researchers noting that citizen science counts may be particularly susceptible to weather induced bias (Dennis *et al*., 2017; Pollard, 1988; Roy *et al*., 2001; Ubach *et al*., 2022). According to the goodness-of-fit analysis, it appears that resident Ringlet and the Common Wood Nymph are more affected by local environmental stochasticity. For these residents, immediate daily weather metrics are of most importance; other studies have shown that specialist butterflies are highly dependent on immediate daily weather for flight activity and detection; they may be alive and present in the area but appear absent during unfavorable weather (Dinsmore *et al*., 2019; Schorr *et al*., 2020). This could be the case for both the ringlet and the Common Wood Nymph.

These micro-scale results complement broader patterns and underscore the importance of investigating community level shifts using this multi-scalar model. Not only were we able to demonstrate that broader climatic shifts facilitate population level resilience, but we were also able to show that finer scale filters influence daily detectability and observed community structure.

## Conclusion

The decline in Shannon Diversity was primarily driven by the severe long-term collapse of the resident, the Common Ringlet; this trend was not universal among all residents as in contrast to the Ringlet, the Common Wood Nymph, another native resident, demonstrated complete long-term stability over the 21-year period. These results indicate that diversity loss in this ecosystem is driven by the asynchronous collapse of a single species rather than a simultaneous decline of all species. The butterfly community in central Alberta has undergone a significant structural shift since 2000 as shown by a decline in the Shannon Diversity index (H). This shows that stable species richness (S) and evenness (J) may mask changes in the community structure and long-term diversity declines. The Ringlet is being affected both on the macro-scale by temporal trends and the micro-scale from immediate weather stressors while having no resilience to buffer population declines. On the other hand, the Cabbage White and Clouded Sulphur are affected mainly by extreme winter minimums. Their multi-voltine flexibility allows them to have boom and bust years but maintain overall long-term population stability allowing them to respond positively to anthropogenic changes. Agricultural fertilization and atmospheric nitrogen deposition have been shown to increase the mortality of common Lepidoptera larvae by at least one-third, challenging the idea that more abundant and generally more widespread species are unaffected by changes in host-plant quality (Kurze et al., 2018). While the more mobile and wider spread species in this Central Alberta community seem to be able to maintain stable populations over time, widespread non-climatic environmental change, such as nitrogen deposition from fertilizer use on farms, represents an unmeasured threat that may prove significant in future years. Given the recent arrival of the European Skipper to the area, we are unsure of what interactions are taking place between it and other species, in particular, the Ringlet. It could also be that the Ringlet is in the process of a climate driven range shift (Breed et al., 2013). This study underpins the necessity to combine multi-decadal survey data with intensive, high-frequency intra-seasonal sampling; this helps to gather information that may prove beneficial in informing conservation priorities that safeguard global butterfly and overall biodiversity. Citizen science, when done in a systematic way, and utilizing standardized methods, is an invaluable resource that has a very powerful role in monitoring and quantifying complex environmental interactions.

## Methods

### Study area and data collection and processing

The transect where butterfly counts were performed is inside a non-for-profit organization located near Lacombe, in Central Alberta, Canada at 52° 28ʹ 06” N, 113° 44ʹ 13” W, elevation 916 m above sea level (Pearman *et al*. 2020). The area is made up of open aspen parkland underlain by sandy soils. The dominant land uses in the area are pastureland for grazing and cropland for crop and forage production (Pearman *et al*. 2006, Merkle and Barclay 1996).

For objectives 1 and 2, we used historical data from single, annual mid-summer counts at the non-for-profit organization during one of their annual events from 2000 to 2025. These counts were started by Charley Bird and performed by the various amateur and professional lepidopterists often associated with the Alberta Lepidopterist Guild (ALG) following similar sampling areas each year. Other notable contributors include Ted Pike, Dave Lawrie, John Acorn, Benny Acorn, along with many others. In some years, surveys consisted of only presence/absence instead of real counts, so those years were eliminated from the analyses. Additionally, for dates between 2021 and 2025 where bug jamboree data was not available, a single count data from intensive multi-day counts was supplemented.

For objective 3, we performed our own, intensive multi-day counts from 2021 - 2025 (exact dates available in supplementary materials). The former Biologist and Site Services Manager, Myrna Pearman, was instrumental in encouraging the starting of these intensive counts. The majority of counts were performed by the second author, with assistance from university students and citizen volunteers who completed training. Efforts were made to conduct counts within 1.5 hours of solar noon to have consistency in our data and because it’s the time of day when butterflies are more active. Deviating slightly from the Pollard protocol (Pollard, 1977), all butterflies identifiable within 5 meters of where the observer was walking were counted; a constant pace was maintained, with occasional stops to record or attempt at catching butterflies that needed a closer examination for identification. Each count lasted approximately 1 hour, and the route taken was always the same; approximately 1 km in length, providing a good variety of habitat for butterflies, and easily accessible (Lewis, 2021; Holtom & Lewis 2022). We obtained a dataset consisting of 77 surveys conducted between 2021 and 2025.

To investigate species-level responses to environmental conditions, we modeled the abundance of the five most dominant taxa in the dataset: Cabbage White (*Pieris rapae),* Clouded Sulphur (*Colias eriphyle),* European Skipper (*Thymelicus lineola),* Ringlet (*Coenonympha tullia)*, and Common Wood Nymph *(Cercyonis pegala)*.

### Weather and Global Climate Metrics

To capture the influence of large-scale climate oscillations, seasonal averages of the Pacific Decadal Oscillation (PDO) and Oceanic Niño Index (ONI) were derived for both winter (December–February) and spring (March–May) periods (https://psl.noaa.gov/data/timeseries/month/). Local environmental stressors were quantified through a Snow Exposure Index, which counts days of extreme cold (below −20°C) paired with minimal snow cover (less than 5 cm), and the Extreme Winter Minimum temperature. Finally, phenological and count-day conditions were controlled for using Spring Growing Degree Days (GDD), the Julian Day of the count, and the specific Temperature for the day of each survey. Survey wind speed was omitted from the multi-decadal analysis due to a high proportion of missing values in the provincial datasets. Historical weather records were obtained from the local LACOMBE CDA 2 station (Station ID: 10906), which provided continuous, uninterrupted coverage for the entire 2000–2025 study period.

### Statistical Analysis: Multi-Model Inference Framework

To evaluate the drivers of butterfly community structure and species-specific abundances across both temporal scales, we employed an Information-Theoretic (IT) multi-model inference framework. Prior to analysis, all continuous weather, climatic, and phenological predictor variables were standardized (Z-scored) to facilitate the direct comparison of effect sizes (Standardized β).

#### a) Multi-Decadal Trends and Climatic Drivers (21-year data)

We fitted global linear models (LM) with a Gaussian error distribution to assess long-term shifts in Shannon Diversity (H) and Pielou’s Evenness (J). For Species Richness (S), which represents discrete count data, we fitted a generalized linear model (GLM) with a Poisson distribution. To evaluate the historical abundance of the five historically dominant taxa, we constructed species-specific global models using Negative Binomial generalized linear models (glm.nb in the MASS package) to appropriately account for the high variance and overdispersion inherent to long-term insect count data. Predictor variables included temporal change (year), local survey temperature, regional overwintering and emergence climate indices (winter_PDO, spring_ONI), and specific local winter/spring weather metrics (snow_exposure_index, extreme_winter_min, spring_gdd).

#### b) Intra-Seasonal High-Resolution Dynamics (2021–2025)

To isolate the fine-scale effects of day-of and lagged weather filters in the 5-year high-resolution dataset, we utilized generalized linear mixed models (GLMMs) via the glmmTMB package. Survey year was included as a random intercept in all short-term models to account for baseline interannual variation. Overdispersed species counts were modeled using a Negative Binomial parameterization (nbinom2). Two specific structural adaptations were required for the high-resolution global models. Because Pielou’s Evenness (J) represents continuous proportion data bounded between 0 and 1, we modeled this index using a GLMM with a Beta error distribution and a logit link; to mathematically accommodate true 1s in the field data (representing even survey days), the response variable was transformed using the Smithson and Verkuilen (2006) scaling formula prior to analysis. Additionally, to accurately model the complex, multivoltine flight phenology of the Clouded Sulphur, a quadratic term for Julian day (Julian Day^2^) was incorporated into the species-specific global model.

#### c) Global Model Validation

Prior to candidate model generation, the structural validity of all overarching global models was rigorously evaluated. To ensure predictor independence, Variance Inflation Factors (VIF) were calculated for all fixed effects to assess multicollinearity, with a threshold of VIF < 3 used to confirm the absence of severe collinearity. Furthermore, simulated residual diagnostics were conducted via the DHARMa package to assess model fit, uniformity, overdispersion, and zero-inflation. Specifically, the inclusion of the quadratic Julian day term for the Clouded Sulphur, and the application of the Smithson transformation paired with a Beta error distribution for Pielou’s Evenness (J), were utilized to resolve non-linear quantile deviations and boundary constraints identified during this diagnostic phase. Only global models that successfully met all underlying distributional assumptions were advanced to the multi-model inference stage.

#### d) Model Selection and Averaging

Candidate models were generated from the global models using the dredge function in the MuMIn package. During the all-subsets evaluation process for the long-term, species-specific datasets, iterative calculation of the Negative Binomial dispersion parameter (ϑ) occasionally resulted in convergence failures for specific mathematically unstable sub-models (notably in the Ringlet and Clouded Sulphur datasets). To resolve this and ensure an uncompromised model selection table, we utilized a fixed-dispersion approach. The successfully converged overdispersion parameter (ϑ) was extracted from the species’ global model. The global model was then refit as a standard generalized linear model (GLM) with the Negative Binomial family utilizing the fixed, pre-calculated ϑ. This stabilized the maximum likelihood estimation across all subsets while preserving the correct variance structure. Furthermore, to prevent multicollinearity during subset selection for the short-term high-resolution dataset, we applied marginality rules to prohibit the simultaneous inclusion of weather metrics on the day of the count and their corresponding 3-day lagged averages within the same candidate model, due to their high correlation.

Candidate models were generated from the global models using the dredge function in the MuMIn package. During the all-subsets evaluation process for the long-term, species-specific datasets, iterative calculation of the Negative Binomial dispersion parameter (ϑ) occasionally resulted in convergence failures for specific mathematically unstable sub-models (notably in the Ringlet and Clouded Sulphur datasets). To resolve this and ensure an uncompromised model selection table, we utilized a fixed-dispersion approach. The successfully converged overdispersion parameter (ϑ) was extracted from the species’ global model. The global model was then refit as a standard generalized linear model (GLM) with the Negative Binomial family utilizing the fixed, pre-calculated ϑ. This stabilized the maximum likelihood estimation across all subsets while preserving the correct variance structure. Furthermore, to prevent multicollinearity during subset selection for the short-term high-resolution dataset, we applied marginality rules to prohibit the simultaneous inclusion of weather metrics on the day of the count and their corresponding 3-day lagged averages within the same candidate model, due to their high correlation. Candidate models were ranked using Akaike’s Information Criterion corrected for small sample sizes (AICc; Burnham & Anderson, 2002). Models with ΔAICc < 2 were considered equally parsimonious and were retained to calculate conditional model-averaged standardized coefficients (β), adjusted standard errors, and 95% Confidence Intervals. The relative variable importance (RVI) for each retained predictor was calculated as the sum of Akaike weights (wi) across all candidate models in the top subset containing that specific term. Finally, to evaluate the goodness-of-fit for the top-ranked models, Nakagawa’s (2013, 2017) marginal and conditional pseudo-R2 values were calculated using the r.squaredGLMM function. All analyses were conducted in R (v4.3.2).

## Author contributions

Both authors contributed equally.

## Code and data availability

The code used in R (v4.3.2) for analyses will be archived in Dryad and made available for peer review during the revision stage. Raw count data for this study are available from the corresponding author upon request.

## Acknowledgments

TD Friends of Environment Grant #75710267 provided funding for training (university students and volunteers), signage, and survey route marking. Naia Holtom assisted in training volunteers, conducting counts and compiling the initial literature review as part of a directed study class. Ellis Nature Centre provided space, historical records, and assisted in volunteer training and Burman University provided seed funds and grant administration. Special thanks to Felix Sperling, John Acorn, and Ainsely Lewis for reviewing the manuscript before submission.

## References

Abarca, M., Larsen, E. A. & Ries, L. (2019) Heatwaves and novel host consumption increase overwinter mortality of an imperiled wetland butterfly. Front. Ecol. Evol. 7, 193.

Au, T. F. & Bonebrake, T. C. (2019) Increased suitability of poleward climate for a tropical butterfly (Euripus nyctelius) accompanies its successful range expansion. J. Insect Sci. 19, 2.

Bollarapu, M. J., Kuchibhotla, S., Kvsn, R. & Patel, H. (2024) Dynamic perspectives on biodiversity quantification: beyond conventional metrics. PeerJ 12, e17924.

Breed, G., Stichter, S. & Crone, E. (2013) Climate-driven changes in northeastern US butterfly communities. Nat. Clim. Change 3, 142–145.

Burnham, K. P., & Anderson, D. R. (2002) Model Selection and Multimodel Inference: A Practical Information-Theoretic Approach (2nd ed.). Springer-Verlag.

Casner, K. L., Forister, M. L., O’Brien, J. M., Thorne, J., Waetjen, D. & Shapiro, A. M. (2014) Contribution of urban expansion and a changing climate to decline of a butterfly fauna. Conserv. Biol. 28, 773–782.

Chen, C., Harvey, J. A., Biere, A. & Gols, R. (2019) Rain downpours affect survival and development of insect herbivores: The specter of climate change? Ecology 100, e02819.

Crossley, M. S., Smith, O. M., Berry, L. L., Phillips-Cosio, R., Glassberg, J., Holman, K. M., Holmquest, J. G., Meier, A. R., Varriano, S. A., McClung, M. R., Moran, M. D., & Snyder, W. E. (2021) Recent climate change is creating hotspots of butterfly increase and decline across North America. Glob. Change Biol. 27, 2702–2714.

de Palma, A., Dennis, R. L. H., Brereton, T., Leather, S. R., & Oliver, T. H. (2016) Large reorganizations in butterfly communities during an extreme weather event. Ecography 39(12), 1189–1197.

Dennis, E. B., Morgan, B. J. T., Brereton, T. M., Roy, D. B. & Fox, R. (2017) Using citizen science butterfly counts to predict species population trends. Conserv. Biol. 31, 1350–1361.

Dinsmore, S. J., Vanausdall, R. A., Murphy, K. T., Kinkead, K. E., & Frese, P. W. (2019) Patterns of monarch site occupancy and dynamics in Iowa. Front. Ecol. Evol. 7, 169.

Dorrestein, A., Rust, H. P. R., Macgregor, N. A., Tiernan, B., Jankowski, A., Woinarski, J. C. Z., James, D. J., Flakus, S., Schulz, M., Pahor, S., Mann, A., Desmond, B., Welbergen, J. A. (2025) Factors affecting the detection probability of a critically endangered flying-fox: consequences for monitoring and conservation. Wildl. Res. 52, WR24030.

Dubey, S., Kalyani, A., Kumar, A. & Chand, P. (2025) Comparative analysis of Simpson’s and Shannon’s biodiversity indices in assessing ecological diversity in Van Vihar region of Jaunpur (U.P.). J. Title 22, 45–47.

Edwards, C. B. et al. (2025) Rapid butterfly declines across the United States during the 21st century. Science 387, 1090–1094.

Flockhart, D. T. T. (2002) The butterfly fauna of Beaverhill Lake, AB. Blue Jay 60, 93–106.

Forister, M. L. et al. (2021) Fewer butterflies seen by community scientists across the warming and drying landscapes of the American West. Science 371, 1042–1045.

Forister, M. L. et al. (2023) Assessing risk for butterflies in the context of climate change, demographic uncertainty, and heterogeneous data sources. Preprint at bioRxiv.

Gezon, Z., Lindborg, R., Savage, A. & Daniels, J. (2018) Drifting phenologies cause reduced seasonality of butterflies in response to increasing temperatures. Insects 9, 174.

Habel, J. C. et al. (2019) Long-term large-scale decline in relative abundances of butterfly and burnet moth species across south-western Germany. Sci. Rep. 9, 14921.

Hallmann, C. A. et al. (2017) More than 75 percent decline over 27 years in total flying insect biomass in protected areas. PLoS One 12, e0185809.

Halsch, C. A., Shapiro, A. M., Forister, M. L. & Grames, E. M. (2025) Shifting baselines in North America’s longest running butterfly monitoring program. Conserv. Lett. 18, e13116.

Hamon, L., Kingsolver, J., Moore, K. & Hurlbert, A. (2024) High voltinism, late-emerging butterflies are sensitive to interannual variation in spring temperature in North Carolina. Environ. Entomol. 54, 77–85.

Hof, A. R. & Svahlin, A. (2016) The potential effect of climate change on the geographical distribution of insect pest species in the Swedish boreal forest. Scand. J. For. Res. 31, 29–39.

Holtom, N. & Lewis, D. (2023) Ellis Bird Farm 2022 Butterfly Count Report. Alberta Lep. Guild Newsl. Spring 2023, 13–16.

Hussain, B., War, A. R., & Pfeiffer, D. G. (2018) Mapping foliage damage index and monitoring of Pieris brassicae by using GIS-GPS technology on cole crops. J. Entomol. Zool. Stud. 6(2), 933–938.

Jones, M. C. et al. (2013) Predicting the impact of climate change on threatened species in UK waters. PLoS One 8, e54216.

Kocher, S. D. & Williams, E. H. (2000) The diversity and abundance of North American butterflies vary with habitat disturbance and geography. J. Biogeogr. 27, 785–794.

Kocsis, M. & Hufnagel, L. (2011) Impacts of climate change on Lepidoptera species and communities. Appl. Ecol. Environ. Res. 9, 43–72.

Konno, K. (2023) Extremely high relative growth rate makes the cabbage white, *Pieris rapae*, a global pest with highly abundant and migratory nature. Sci. Rep. 13, 9697.

Kurze, S., Heinken, T. & Fartmann, T. (2018) Nitrogen enrichment in host plants increases the mortality of common Lepidoptera species. Oecologia 188, 1227–1237.

Kuussaari, M. et al. (2016) Weather explains high annual variation in butterfly dispersal. Proc. R. Soc. B, 283(1834), 20160413.

Kuussaari, M. et al. (2021) Butterfly species’ responses to urbanization: differing effects of human population density and built-up area. Urban Ecosyst. 24, 515–527.

Larsen, E. A. et al. (2024) Overwintering strategy regulates phenological sensitivity with consequences for ecological services in a clade of temperate North American insects. Funct. Ecol. 38, 1075–1088.

Leston, L., & Koper, N. (2016) Urban rights-of-way as extensive butterfly habitats: A case study from Winnipeg. Landsc. Urban Plan. 157, 56–62.

Leston, L., & Koper, N. (2019) An urban wildlife habitat experiment: conservation implications of altering management regimes on animals and plants along urban and rural rights-of-way. J. Urban Ecol. 5(1), juz013.

Leuenberger, W. et al. (2025) Three decades of declines restructure butterfly communities in the Midwestern United States, Proc. Natl. Acad. Sci. 122 (33), e2501340122.

Lewis, D. (2022) Ellis Bird Farm 2021 Butterfly Count Report. Alberta Lep. Guild Newsl. Spring 2022, 18–22.

Li, Y. et al. (2023) Extinction risk modeling predicts range-wide differences of climate change impact on Karner blue butterfly (*Lycaeides melissa samuelis*). PLoS One, 18(11), e0262382.

MacDonald, Z. G. et al. (2017) Negative relationships between species richness and evenness render common diversity indices inadequate for assessing long-term trends in butterfly diversity. Biodivers. Conserv. 26, 617–629.

McDermott Long, O., et al. (2017) Sensitivity of UK butterflies to local climatic extremes: which life stages are most at risk? J. Anim. Ecol. 86(1), 108–116.

Mena, S., Rentería, J., & Checa, M. F. (2025) Diel Versus Seasonal Butterfly Community Partitioning in a Hyperdiverse Tropical Rainforest. Insects 16(12), 1247.

Nakagawa, S., & Schielzeth, H. (2013) A general and simple method for obtaining R^2 from generalized linear mixed-effects models. Methods Ecol. Evol. 4(2), 133–142.

Nakagawa, S., Johnson, P. C. D., & Schielzeth, H. (2017) The coefficient of determination R^2 and intra-class correlation coefficient from generalized linear mixed-effects models revisited and expanded. J. R. Soc. Interface 14(134), 20170213.

Nowicki, P., Settele, J., Henry, P. & Woyciechowski, M. (2008) Butterfly monitoring methods: the ideal and the real world. Isr. J. Ecol. Evol. 54, 69–88.

Pilliod, D. S. & Rohde, A. T. (2016) Insect community responses to climate and weather across elevation gradients in the Sagebrush Steppe, eastern Oregon. U.S. Geological Survey.

Pinkert et al. (2025) Global hotspots of butterfly diversity are threatened in a warming world. Nat. Ecol. Evol, 9(5), 789–800.

Pollard, E. (1977) A method for assessing changes in the abundance of butterflies. Biol. Conserv. 12, 115–134.

Pollard, E. (1988) Temperature, rainfall and butterfly numbers. J. Appl. Ecol. 25(3), 819–828.

Prudic, K. L. et al. (2017) eButterfly: leveraging massive online citizen science for butterfly conservation. Insects 8, 53.

Ragonese, I. G. et al. (2024) Extreme heat reduces host and parasite performance in a butterfly–parasite interaction. Proc. R. Soc. B 291, 20232305.

Ries, L. & Oberhauser, K. (2015) A citizen army for science: quantifying the contributions of citizen scientists to our understanding of monarch butterfly biology. BioScience 65, 419–430.

Riva, F., Acorn, J. H. & Nielsen, S. E. (2018) Localized disturbances from oil sands developments increase butterfly diversity and abundance in Alberta’s boreal forests. Biol. Conserv. 217, 173–180.

Robbins, C. S. (1981) Bird activity levels related to weather. Stud. Avian Biol. 6, 301–310.

Roland, J. & Matter, S. F. (2013) Variability in winter climate and winter extremes reduces population growth of an alpine butterfly. Ecology 94, 190–199.

Roland, J. & Matter, S. F. (2016) Pivotal effect of early-winter temperatures and snowfall on population growth of alpine Parnassius smintheus butterflies. Ecol. Monogr. 86, 412–428.

Roy, D. B. et al. (2001) Butterfly numbers and weather: Predicting historical trends in abundance and the future effects of climate change. J. Anim. Ecol. 70(2), 201–217.

Ryan, S. F. et al. (2019) Global invasion history of the agricultural pest butterfly *Pieris rapae* revealed with genomics and citizen science. Proc. Natl Acad. Sci. USA 116, 20015–20024.

Sattar, Q., Maqbool, M. E., Ehsan, R. & Akhtar, S. (2021) Review on climate change and its effect on wildlife and ecosystem. Open J. Environ. Biol. 6, 008–014.

Schorr, R. A. et al. (2020) Multi-year occupancy of the hops blue butterfly (*Celastrina humulus*): habitat patch colonization and extinction. J. Insect Conserv. 24, 927–934.

Sevilleja, C. G. et al. (2020) Assessing Butterflies in Europe – European Butterfly Monitoring Scheme - Network development: Technical Report. Butterfly Conservation Europe.

Smithson, M., & Verkuilen, J. (2006) A better lemon squeezer? Maximum-likelihood regression with beta-distributed dependent variables. Psychol. Methods 11(1), 54–71.

Teichman, K. J., Nielsen, S. E. & Roland, J. (2013) Trophic cascades: linking ungulates to shrub-dependent birds and butterflies. J. Anim. Ecol. 82, 1288–1299.

Ubach, A. et al. (2022) Weather and butterfly responses: A framework for understanding population dynamics in terms of species’ life-cycles and extreme climatic events. Oecologia, 199(2), 243–259.

Ulrich, W. et al. (2016) Environmental correlates of species rank - abundance distributions in global drylands, Perspect. Plant Ecol. Evol. Syst. 20, 56–64.

Van Strien, A. J. et al. (2017) Opportunistic citizen science data of butterflies and dragonflies: the importance of quantifying sampling effort. Conserv. Biol. 31, 1350–1361.

Vitasse, Y. et al. (2021) Phenological and elevational shifts of plants, animals and fungi under climate change in the European Alps. Biol. Rev. 96, 1816–1835.

von Schmalensee, L. et al. (2023) Seasonal specialization drives divergent population dynamics in two closely related butterflies. Nat Commun. 14(1), 3663.

Wepprich, T. et al. (2019) Butterfly abundance declines over 20 years of systematic monitoring in Ohio, USA. PLoS One 14, e0216270.

Wikström, L., Milberg, P., & Bergman, K.-O. (2009) Monitoring of butterflies in semi-natural grasslands: Diurnal variation and weather effects. J. Insect Conserv. 13(2), 203–211.

